# Long-read RNA Sequencing Improves the Annotation of the Equine Transcriptome

**DOI:** 10.1101/2022.06.07.495038

**Authors:** S. Peng, AR. Dahlgren, EN. Hales, A.M. Barber, T. Kalbfleisch, JL. Petersen, RR. Bellone, M. Mackowski, K. Cappelli, S. Capomaccio, SJ. Coleman, O. Distl, E. Giulotto, B. Waud, N.A. Hamilton, T. Leeb, G. Lindgren, LA. Lyons, M. McCue, JN. MacLeod, J. Metzger, JR. Mickelson, BA. Murphy, L. Orlando, C. Penedo, T. Raudsepp, E. Strand, T. Tozaki, DS. Trachsel, BD. Velie, CM. Wade, J. Cieslak, CJ. Finno

## Abstract

A high-quality reference genome assembly, a biobank of diverse equine tissues from the Functional Annotation of the Animal Genome (FAANG) initiative, and incorporation of long-read sequencing technologies, have enabled efforts to build a comprehensive and tissue-specific equine transcriptome. The equine FAANG transcriptome reported here provides up to 45% improvement in transcriptome completeness across tissue types when compared to either RefSeq or Ensembl transcriptomes. This transcriptome also provides major improvements in the identification of alternatively spliced isoforms, novel noncoding genes, and 3’ transcription termination site (TTS) annotations. The equine FAANG transcriptome will empower future functional studies of important equine traits while providing future opportunities to identify allele-specific expression and differentially expressed genes across tissues.

## 2 Introduction

Equine genome assemblies^1,2^ have provided vital resources for equine genetics research allowing for tools to be built. However, it is evident that detailed annotation of the genome is necessary for further investigation of both simple and complex traits in horses. Current equine genome assemblies have annotations provided by both Ensembl^3,4^ and NCBI^5,6^ gene annotation pipelines. These annotations relied primarily on limited available RNA-seq data, cross-species alignments, and computational predictions. The equine RefSeq annotation release 103 from NCBI contains 33,146 genes and pseudogenes, of which 21,129 are protein coding, and 77,102 transcripts, including 60,887 mRNAs and 16,215 non-coding RNAs^6^. This presents a total isoform-to-gene ratio of 2.3, or 2.8 if only coding genes are considered. Similarly, the equine Ensembl annotation release 105.3 contains 20,955 protein coding genes and 9,014 non-coding genes, with 59,087 transcripts, resulting in a transcript-to-gene ratio of 2.0^4^. For comparison, the most recent GENCODE human gene annotation release (release 39, GRCh38.p13) includes 61,533 genes, of which 19,982 are protein coding, with an average isoform-to-gene ratio of 3.9, or 4.3 when only considering protein coding genes^7^. This presents the question of potentially missing alternatively spliced isoforms in the current equine gene annotation. The human ENCODE projects determined that genes tend to express many isoforms simultaneously but different dominant isoforms exist in different cell lines^8^. The tissue-specific nature of isoform expression underscores the urgent need for a transcriptome with more complete isoform annotation.

Additionally, noncoding RNAs play an important role in many biological pathways^9–12^. With the rising popularity of noncoding RNA therapeutics^13^, a comprehensive noncoding RNA annotation for the horse genome will certainly be an asset to the equine research community. The Ensembl annotation for EquCab3 includes 9,014 noncoding genes while RefSeq annotation contains 8,893 noncoding genes. In comparison, GENCODE human gene annotation release 39 includes 26,378 noncoding genes. A particular challenge with annotating noncoding RNAs comes from the fact that noncoding RNAs are usually less evolutionarily conserved^14^ and present at very low levels^15,16^. Therefore, without deep sequencing of diverse tissue types, noncoding RNAs typically remain unannotated. Since the current equine gene annotation relies heavily on cross-species conservation, and a limited number of RNA-seq data that are publicly available, it is expected that a large number of noncoding RNAs are currently unannotated.

To address these challenges, the equine Functional Annotation of Animal Genome (FAANG) project has collected over 80 tissue types and body fluids from 4 adult Thoroughbred horses (two females and two males)^17,18^. These horses underwent thorough clinical examinations and were selected as healthy references. The FAANG biobank has produced a diverse dataset describing various aspects of the equine gene regulation^19–21^. Here, we report our efforts to build a comprehensive transcriptome for the horse genome using long-read sequence technologies across an array of diverse tissues.

## 3 Results

### 3.1 Transcript Annotation

Full-length non-redundant transcripts were categorized based on their annotated splice junctions compared to reference Ensembl transcripts^4,22^ and the genomic overlap between the two, following the schematics introduced by Tardaguila *et al*^23^. Overall, isoforms with novel splice sites (novel not in catalog, NNC) account for over 40% of all Iso-seq transcripts (**Table 1**), highlighting the insufficiency of short-read data for splice junction discoveries. The majority of novel genes identified in the Iso-seq transcriptome have only one isoform per gene, which are predominantly mono-exonic (**Fig 1A-D**). Compared to the Ensembl annotation, the Iso-seq transcriptome contains fewer short transcripts (<0.5 Kb) but substantially more long transcripts (>1.5 Kb) (**Fig 1E**).

**Table 1.** FAANG transcripts breakdown by structural category. FSM: full-splice match; ISM: incomplete-splice match; NIC: novel-in-catalog; NNC: novel-not-in-catalog

| **Structural Category** | **Count** | **Percentage** |
| --- | --- | --- |
| Antisense | 928 | 1.64 |
| FSM | 12470 | 22.01 |
| Fusion | 684 | 1.21 |
| Genic | 2137 | 3.77 |
| ISM | 4937 | 8.71 |
| Intergenic | 4543 | 8.02 |
| NIC | 6330 | 11.17 |
| NNIC | 24634 | 43.47 |

**Figure 1.**
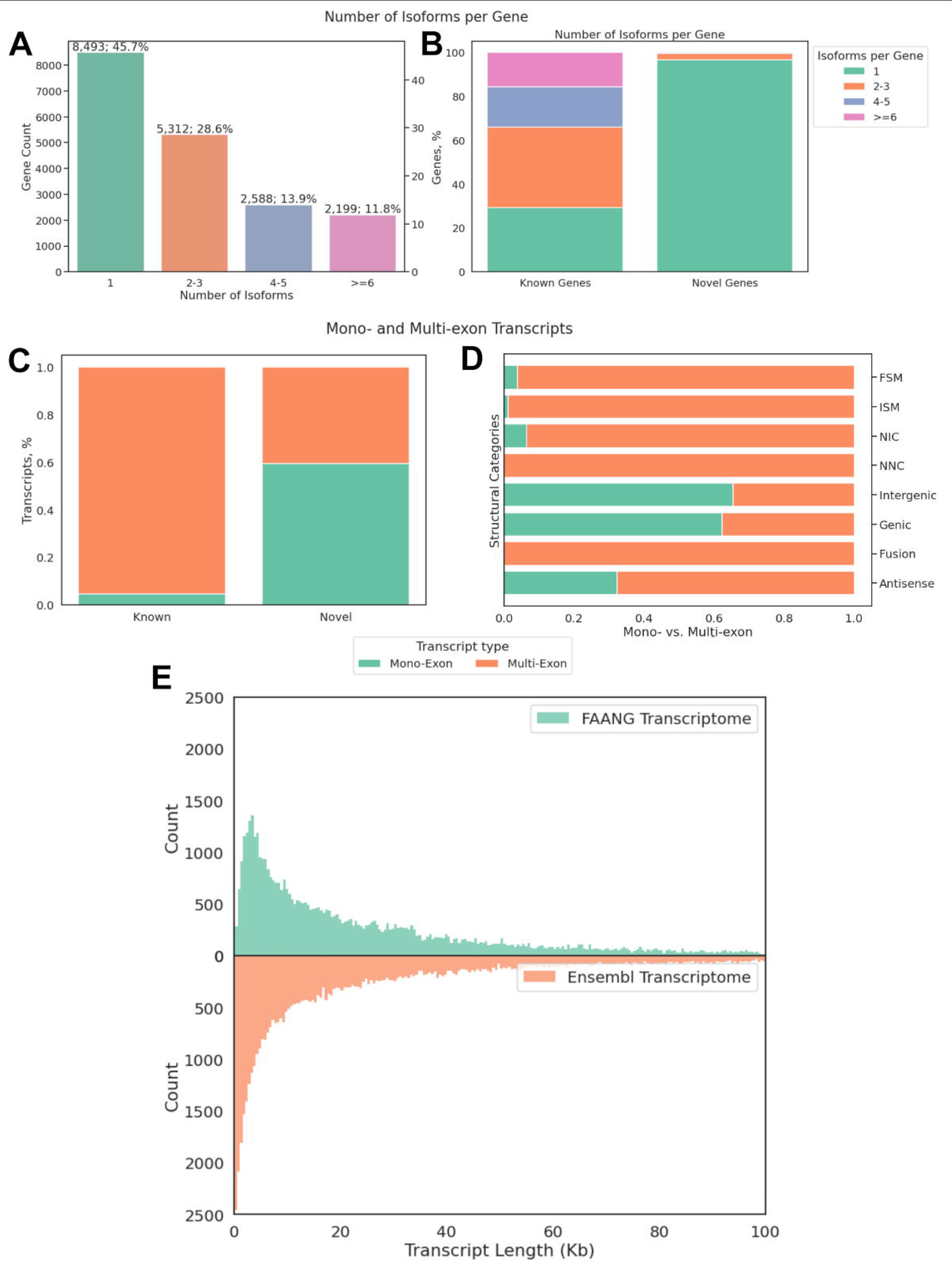
Summary of the FAANG equine transcriptome. The overall number of isoforms (A), known vs. novel genes (B), percentages of known and novel transcripts (C) and portion of mono- and multi-exon transcripts by structural categories that were annotated in the FAANG equine transcriptome. (D) Percentages of mono- and multi-exonic transcripts in each structural category; FSM: full-splice match, all exons and splice junctions match a known reference; ISM: incomplete-splice match, like FSM but missing 3’ and/or 5’ ends; NIC: novel-in-catalog, novel transcripts with known exons and splicing sites; NNC: novel-not-in-catalog, novel transcripts with novel splicing sites; Intergenic: novel transcripts with no overlapping known genes; Genic: novel transcripts overlapping known introns; Fusion: fusion transcripts; Antisense: novel transcripts on the opposite strand of known transcripts (E) The FAANG transcriptome contained fewer short transcripts (<0.5 Kb) but a greater number of long transcripts (>1.5 Kb) as compared to the Ensembl transcriptome.

### 3.2 5’ Completeness

Since standard Iso-seq libraries do not capture 5’ caps of transcripts^24^, we used an aggressive collapsing approach where all transcripts with identical 3’ ends and only differing at 5’ ends are merged into a single transcript. While this approach may hinder the discovery of alternative transcription start sites (TSS), it was necessary to prevent 5’ degraded transcripts from being falsely identified as unique isoforms. To assess the completeness of 5’ ends of annotated transcripts, short-read RNA-seq and ATAC-seq data from the same tissues were used to compare coverage near annotated TSS. Overall, 98.4% transcripts have higher RNA-seq coverage in the 100bp window downstream of the Iso-seq annotated TSS than upstream (**Fig 2A**). Transcripts with a log_2_ ratio of greater than 1 were designated as 5’ complete. A majority of transcripts across all structural categories were determined to have complete 5’ ends, with novel genes (genic, intergenic, and antisense) having a greater percentage of 5’ incomplete transcripts (**Fig 2B**). Additionally, ATAC-seq of the same tissues also show substantial enrichment at annotated TSS (**Fig 2C**).

**Figure 2.**
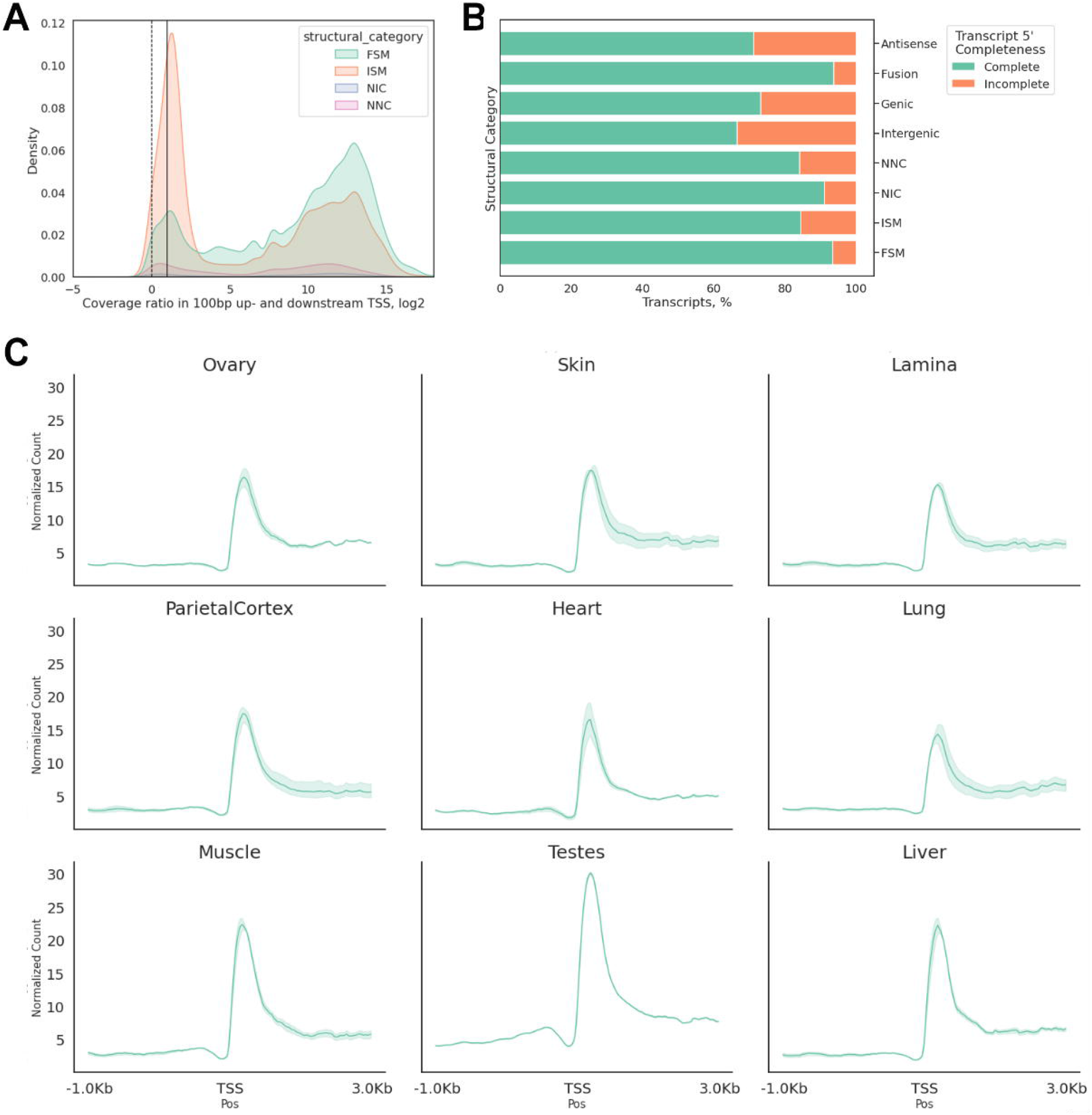
5’ Completeness of the FAANG equine transcriptome. (A) Log_2_ of 100 bp downstream over upstream of TSS RNA-seq coverage. Positive ratios indicate higher coverage downstream of TSS. Dotted line indicates equal coverage up- and down-stream of TSS; solid line indicates 100% higher coverage down-stream TSS than up-stream (B) Percentages of transcripts whose log_2_ ratios are greater than 1, denoted as 5’ complete (green) (C) ATAC-seq read coverage in 1 kb upstream and 3 kb downstream of annotated TSS. FSM: full-splice match; ISM: incomplete-splice match; NIC: novel-in-catalog; NNC: novel-not-in-catalog; Schematics defined by Tardaguila et al^23^.

### 3.3 3’ Completeness

To capture polyadenylated transcripts with complete 3’ ends, poly-T oligonucleotides were used during library construction of Iso-seq. However, internal stretches of adenines could also bind to poly-T oligonucleotides, a phenomenon known as intra-priming, which results in truncated transcripts^23^. To detect potential intra-primed transcripts, the percentage of adenines in a 20 bp window immediately downstream of the annotated transcription termination site (TTS) was calculated for every Iso-seq transcript. Transcripts with 80% or more adenines (i.e., allowing for 4 mismatches with poly-T oligonucleotides) in a 20 bp window downstream of annotated TTS were designated as potential intra-priming candidates. Multi-exonic transcripts across all structural categories had approximately 25% adenines on average, with fewer than 5% transcripts having over 80% adenines, suggesting a high level of 3’ completeness (**Fig 3A**). Over 30% of mono-exonic isoforms with novel exons (NIC) are flagged as potentially intra-primed (**Fig 3B**). Many of these transcripts retain a partial intron and may be intron-retaining isoforms undergoing nonsense-mediated decay (NMD). A comparison between Iso-seq and Ensembl annotation showed significant improvement of TTS, as well as minor improvements of TSS annotation for full-splice matched (FSM) transcripts (**Fig 3C**).

**Figure 3.**
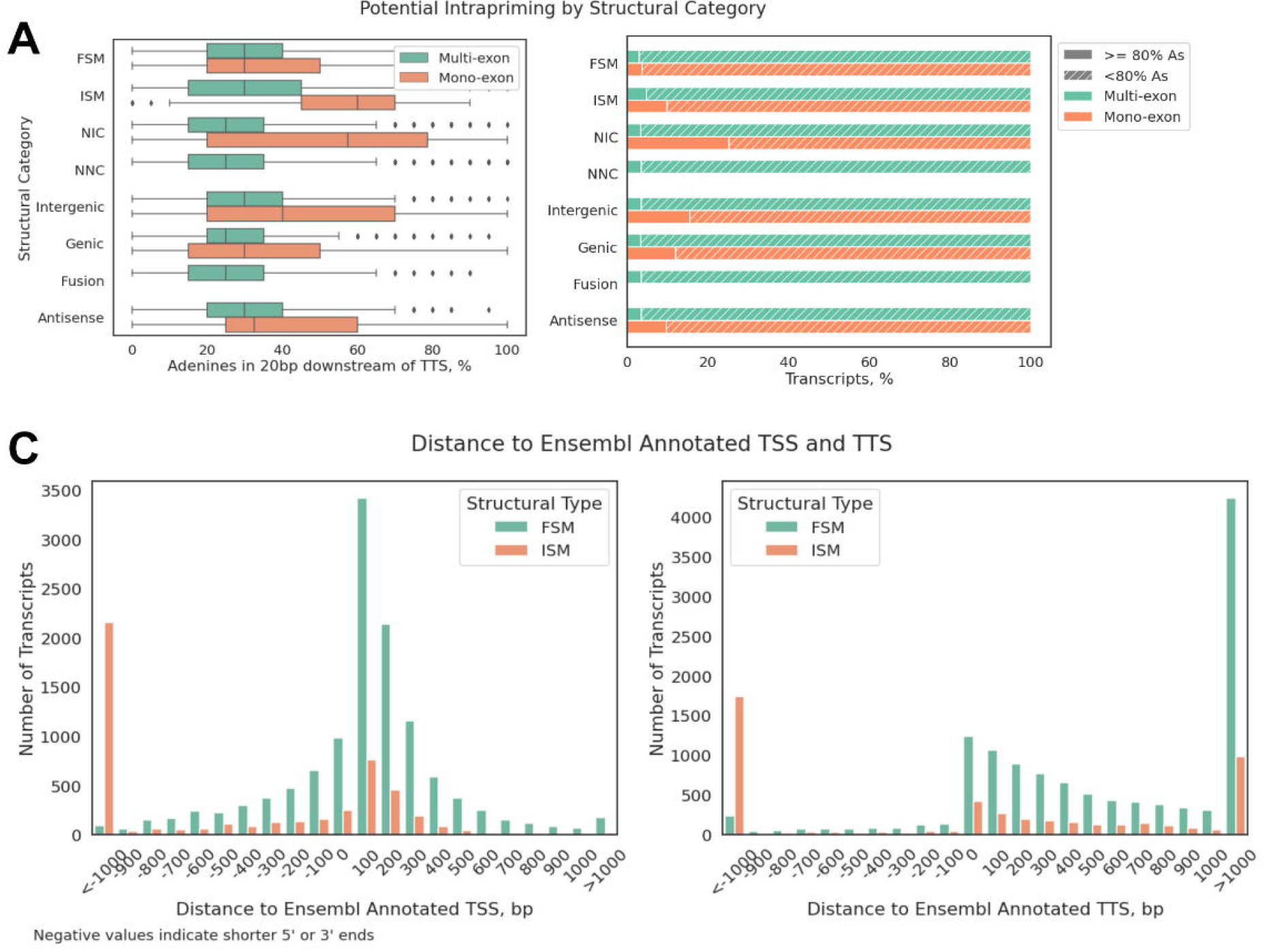
3’ Completeness of the FAANG equine transcriptome. (A) Percentages of adenines in 20 bp genomic regions immediately downstream of annotated TTS. Boxes indicate interquartile range (IQR) and whiskers indicate 1.5*IQR (B) Portions of transcripts with more than 80% adenines in 20bp genomic regions immediately downstream of annotated TTS, by structural categories and exon counts. (C) Distance between FAANG annotated TTS/TSS and Ensembl annotated TTS/TSS, negative values indicate shorter 5’ or 3’ ends. Eg. -1000 indicates that FAANG annotated TSS is 1000 bp downstream of Ensembl annotated TSS (left) or that FAANG annotated TTS is 1000 bp upstream of Ensembl annotated TTS (right).

### 3.4 Protein Coding and Noncoding Transcripts

Open reading frames (ORFs) were predicted using GeneMarkS-T (GMST) algorithm^25^ by SQANTI3^23^ to identify protein-coding transcripts in the FAANG transcriptome. The vast majority of transcripts belonging to known genes had ORFs (97.6% of FSM, 96.8% of ISM, 92.5% of NIC, and 95.4% of NNC), while a significant proportion of novel genes had transcripts without ORFs (28% of genic, 60% of intergenic, and 67.6% antisense transcripts) (**Fig 4A**). There is also a substantial difference in exon counts among coding and noncoding transcripts, with 44.6% of noncoding transcripts being mono-exonic as compared to 4.6% of coding transcripts. Specifically, coding transcripts with novel junctions (NIC) are 96.8% multi-exonic while noncoding NIC transcripts are 56% multi-exonic. Similarly, coding transcripts that overlap or fall within annotated introns are 53% multi-exonic, while only 7.5% of those without an ORF are multi-exonic (**Fig 4B**).

**Figure 4.**
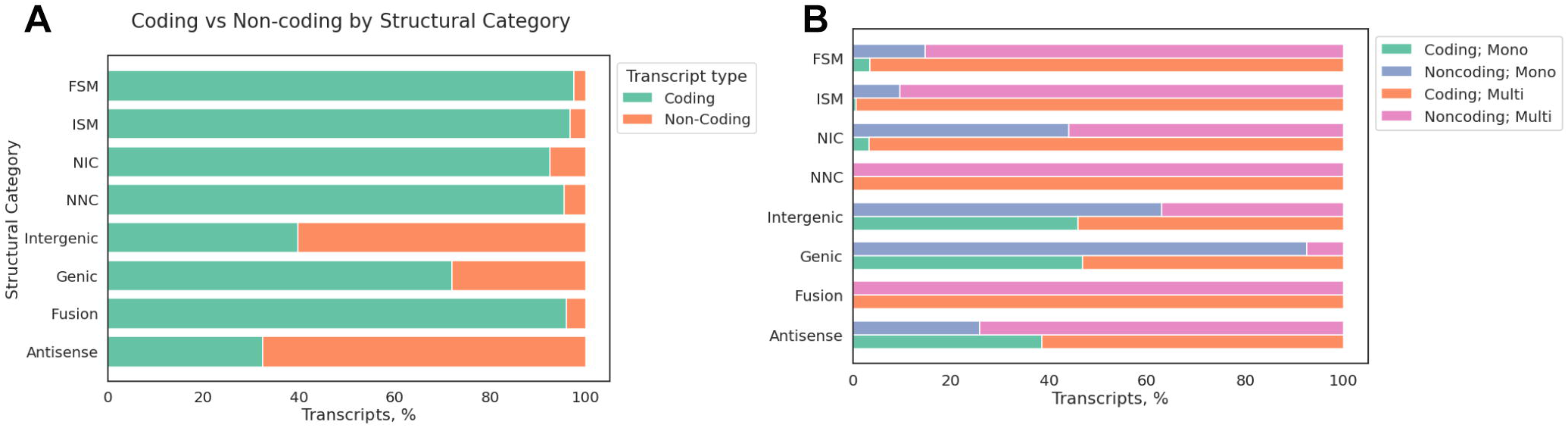
Protein coding and non-coding transcripts in the FAANG equine transcriptome. (A) Portions of coding vs. noncoding transcripts by structural categories and (B) portions of coding vs. noncoding transcripts by structural categories and exon counts.

### 3.5 Splice Junctions

Any junctions not covered by at least one uniquely mapped read from RNA-seq data were removed, along with their associated transcripts. A total of 8,476 transcripts containing 14,738 such junctions were removed at this step. Known junctions on average had 4.8x RNA-seq coverage as compared to novel junctions. This difference primarily came from canonical junctions (GT-AG, GC-AG and AT-AC) (**Fig 5A**). Novel isoforms and transcripts of novel genes also have lower minimum junction coverage as compared to known isoforms (FSM and ISM, Kruskal-Wallis H-test, p<0.0001; post-hoc Dunn’s test p <3.5 × 10^−68^, Bonferroni corrected α=0.003**; Fig 5B**).

**Figure 5.**
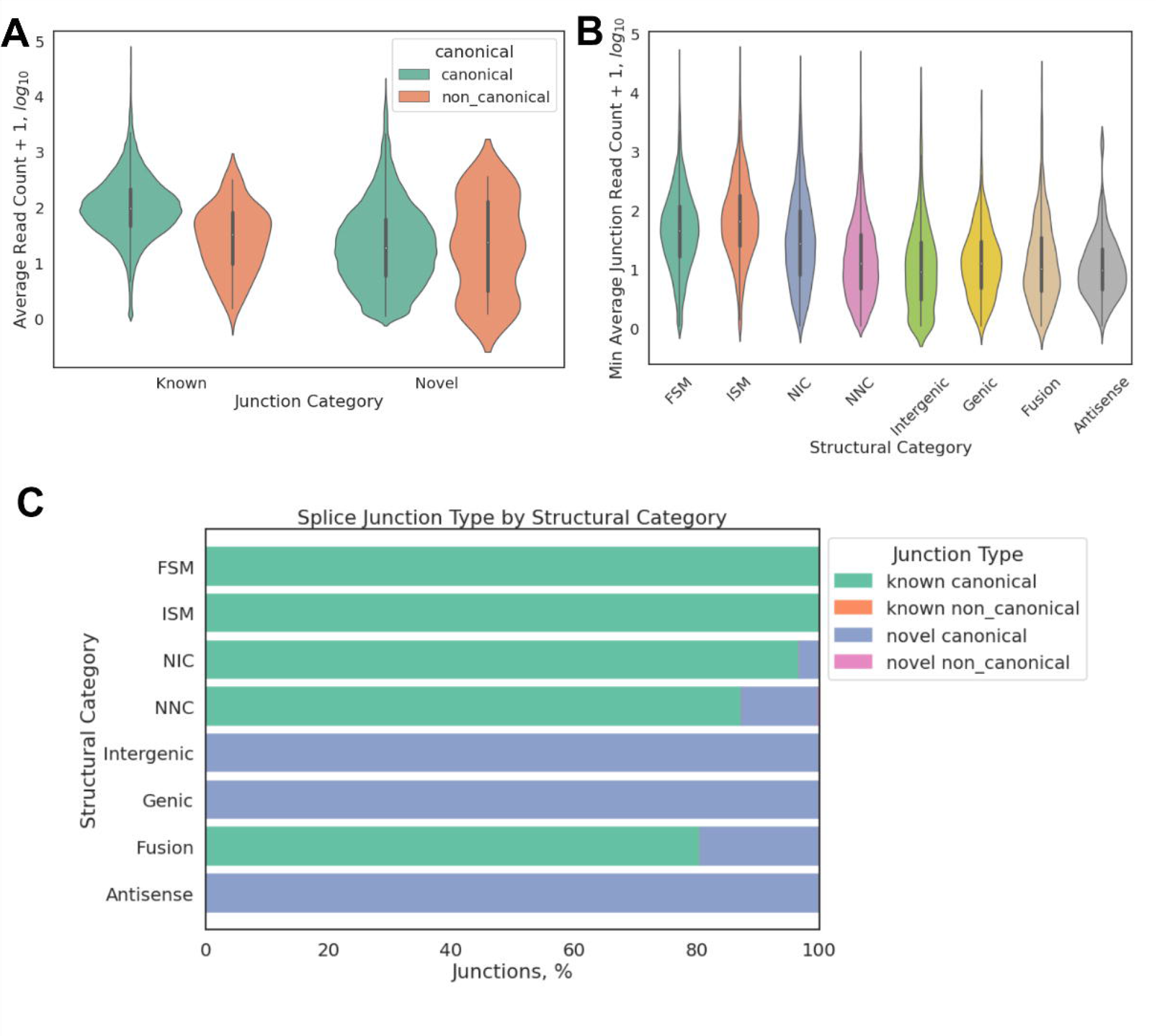
Splice junctions are better defined in the FAANG equine transcriptome. (A) RNA-seq coverage measured as log_10_(ReadCount+1) at known or novel splice junctions, (B) minimum splice junction coverage transcripts by structural categories and (C) splice junction types by structural categories, known non-canonical junctions were observed in FSM, ISM, NIC, and NNC at 2.2%, 0.2%, 2.0%, and 1.2%, respectively; novel non-canonical junctions were only observed in NNC and intergenic isoforms at 2.8% and 2.5%, respectively.

About 10% of the splice junctions annotated in Iso-seq transcriptome were novel compared to Ensembl transcriptome (56,503 out of 581,782). These novel splice junctions contributed discoveries of 36,795 novel isoforms (**Fig 5C**). The GT-AG splice site is observed in 99.2% of splice junctions, with GC-AG and AT-AC sites observed in 0.68% and 0.05% of transcripts, respectively. Non-canonical splice sites were primarily observed at very low frequencies (<3%) (**Fig 5C**; **Table 2**).

**Table 2.**
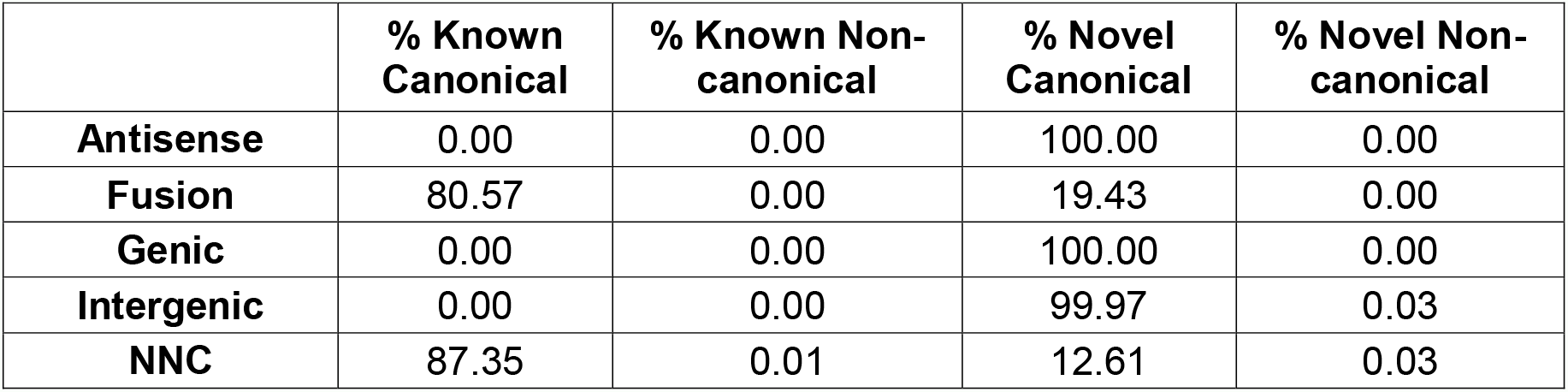
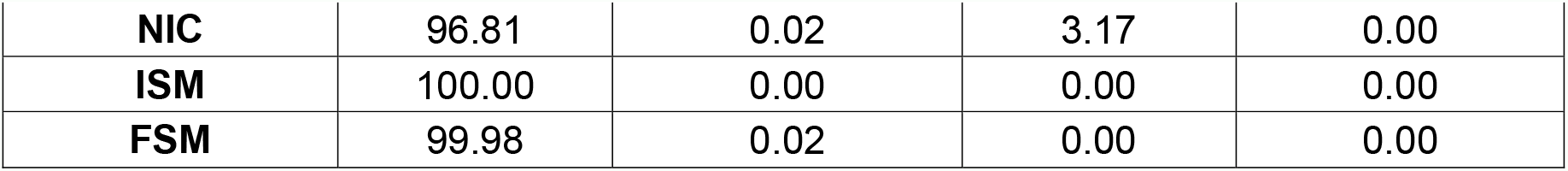
Portions of different splice junction types by structural categories. FSM: full-splice match; ISM: incomplete-splice match; NIC: novel-in-catalog; NNC: novel-not-in-catalog

### 3.6 Sense-Antisense Transcripts

There were a total of 861 novel antisense transcripts identified, with 2,742 isoforms annotated on the opposite strand. Overall, we identified 3,246 transcripts on the plus strand that overlap at least 1 bp with a transcript on the minus strand. Among these sense-antisense pairs of transcripts, 2,249 (69.3%) were coding-to-coding pairs, 954 (29.4%) coding-to-noncoding pairs, and 43 (1.3%) noncoding-to-noncoding pairs.

### 3.7 Tissue-specific Expression

Short-read RNA-seq data from 57 tissues (46 tissues from female animals and 23 tissues from male animals, with 12 tissues from both sexes, **Supplementary Table 1**) were used to quantify the Iso-seq transcriptome. Most known isoforms were ubiquitously expressed in the majority of tissues sequenced, while novel isoforms of known genes and novel intergenic transcripts each showed a bimodal distribution, with many transcripts detected in only a small number of tissues (**Fig 6A**). We also noted that the majority of multi-isoform genes expressed more than one isoform in any given tissue (**Fig 6B**) and had different dominant major isoforms (isoform with highest relative expression of a given gene), depending on the tissue type (**Fig 6C-D**). Similar to humans, major isoforms in horses usually account for 30%-70% of corresponding genes’ total expression in any given tissue (**Fig 6E**).

**Figure 6.**
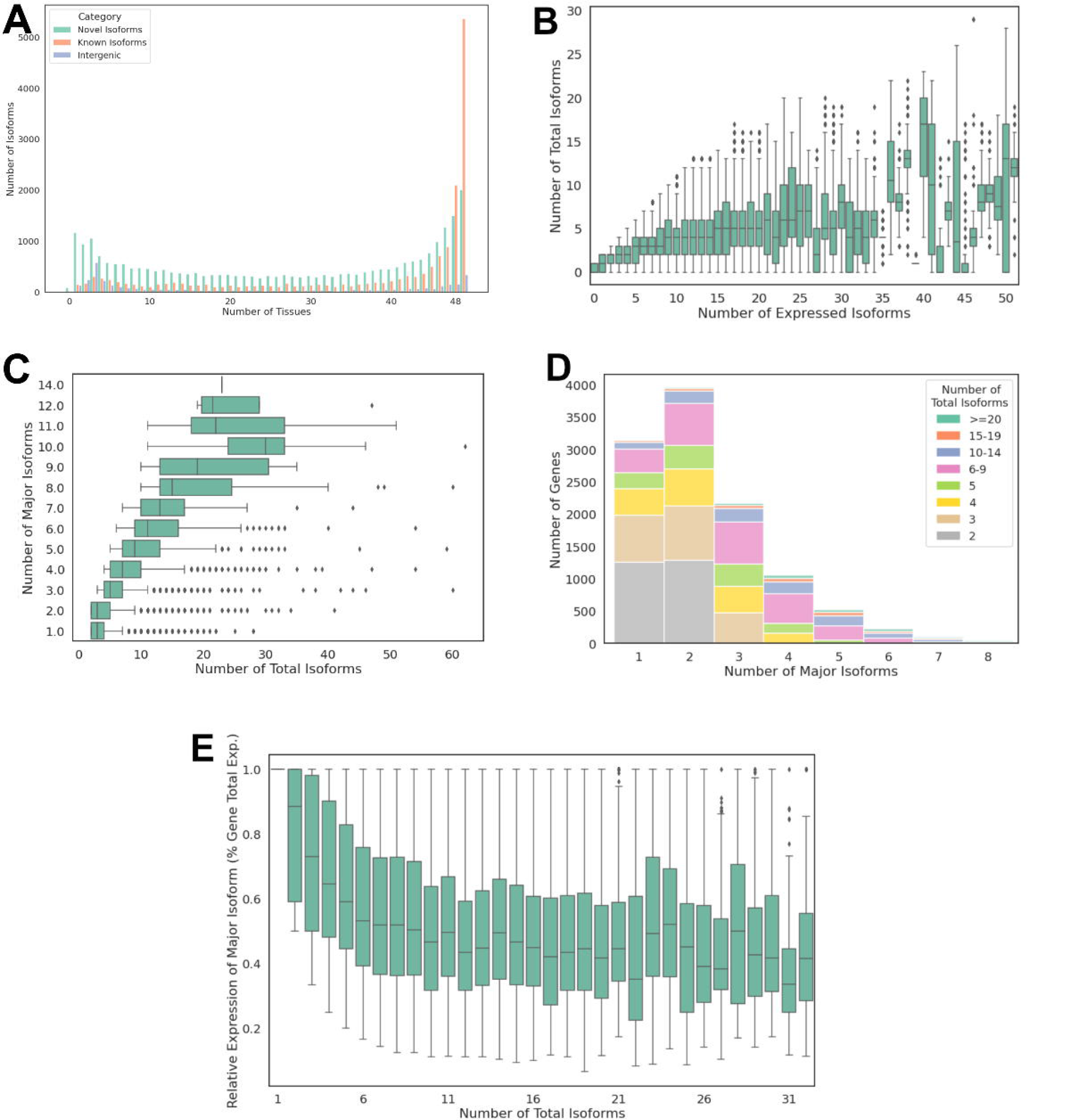
Short-read RNA-seq data mapped to the FAANG equine transcriptome identifies tissue-specific isoforms. (A) Distribution of known vs. novel transcripts detected in different numbers of tissues, (B) number of expressed isoforms across all tissues vs. number of total isoforms per gene; boxes indicate IQR and whiskers indicate 1.5*IQR, (C) number of different major isoforms expressed across tissues vs. number of total isoforms annotated, (D) distribution of genes with different number of major isoforms and their total annotated isoforms and (E) relative expression of major isoforms in each tissue vs. total number of isoforms annotated.

Notably, our RNA-seq data exhibited prominent clustering between sexes within the CNS tissues (**Fig 7A**). Since mare and stallion tissues were prepared at two different laboratories, despite using same protocols, it is unclear whether this clustering reflected genuine biological differences between sexes or was a result of batch effects arising from the RNA isolations. Nonetheless, we observed expected clustering of the other major tissue types across all samples (**Fig 7B**).

**Figure 7.**
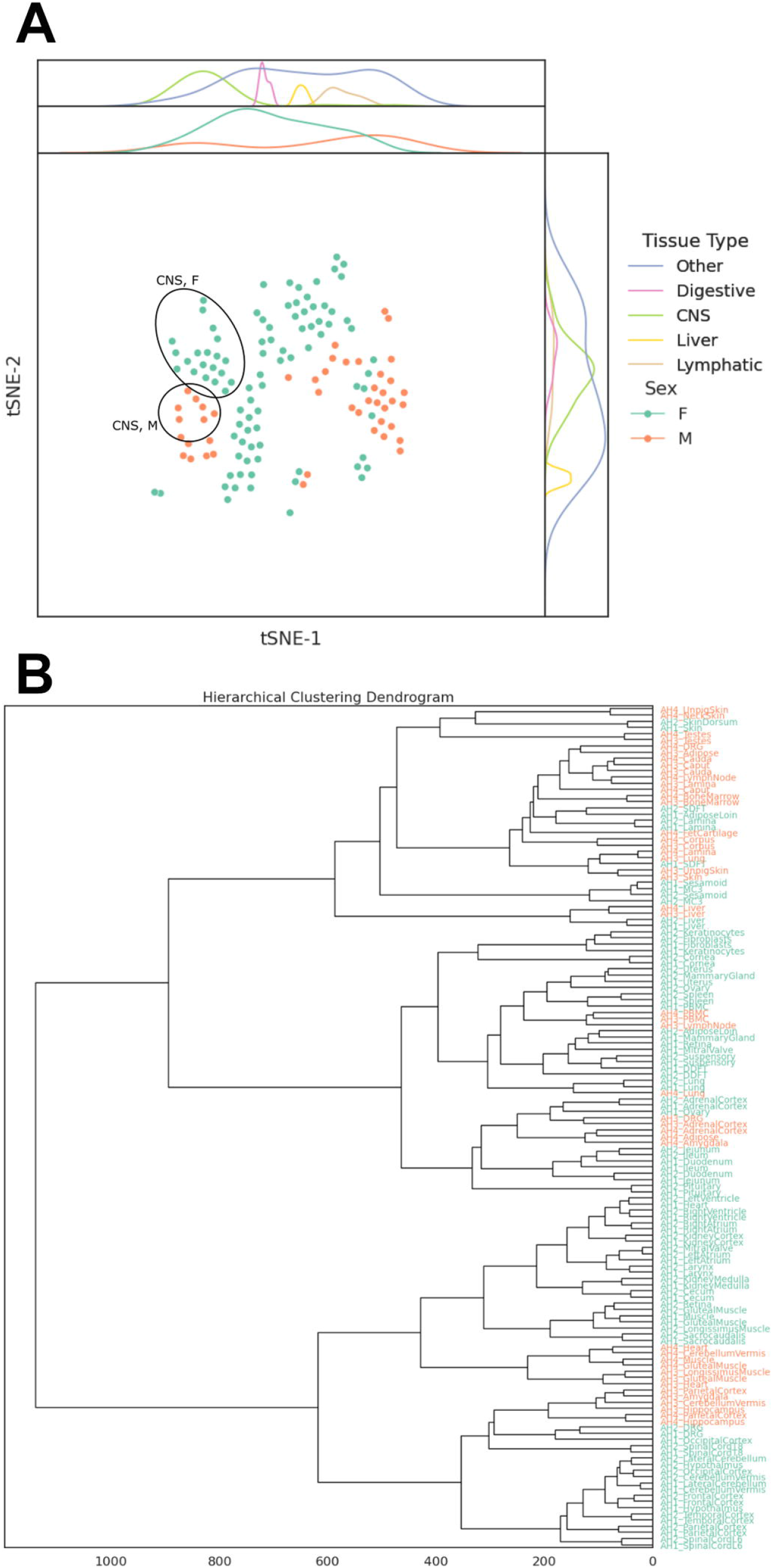
Sex-specific clustering of gene expression across tissue types. (A) t-SNE plot of transcript levels (TPM) across samples and tissues and (B) agglomerative clustering of RNA-seq samples.

### 3.8 Comparison with Ensembl and RefSeq Transcriptomes

To assess the completeness of the Iso-seq transcriptome, RNA-seq reads were aligned to the reference genome and portions of the reads that overlapped a transcribed region annotated in the Iso-seq, Ensembl, or RefSeq transcriptome were calculated as fraction of reads in transcript (FRiT). Considerable improvement in FRiT was observed in tissues that were directly utilized to generate the Iso-seq transcriptome (**Fig 8A**). Finally, we aligned all available RNA-seq reads directly to each transcriptome and observed 3-23% improvement in numbers of properly paired reads in all tissues but cerebellum vermis, duodenum, fibroblasts, keratinocytes, bone marrow, and epididymis (caput, corpus, and cauda) (**Fig 8 B**).

**Figure 8.**
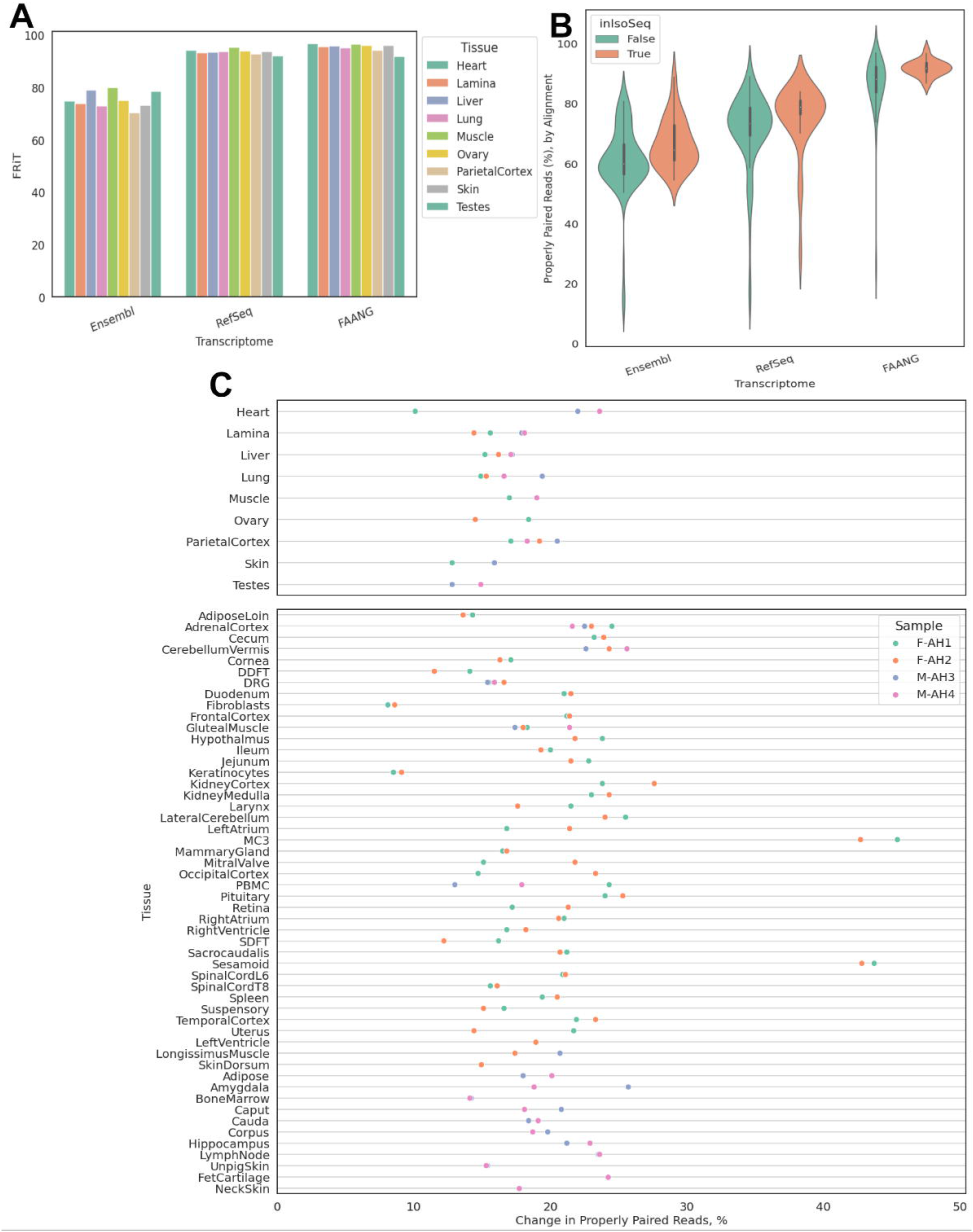
Comparison of FAANG, RefSeq and Ensembl equine transcriptomes. (A) Fractions of reads in transcriptome (FRiT) from RNA-seq reads against FAANG, RefSeq, or Ensembl transcriptomes, (B) distribution of percentages of properly paired reads when RNA-seq data is aligned directly to FAANG, RefSeq, or Ensembl transcriptomes and (C) changes in percentages of properly paired reads aligned to combined FAANG transcriptome when compared to Ensembl or RefSeq transcriptomes, whichever has higher percentage.

To provide a comprehensive set of transcripts for the equine genome, we combined the Iso-seq transcriptome with Ensembl and RefSeq transcriptomes into a single annotation, termed the equine FAANG transcriptome. The FAANG transcriptome consists of 153,492 transcripts (of which 128,723 are multi-exonic) from 36,239 genes, with a gene-to-isoform ratio of 4.2. This combined transcriptome contains a total of 26,631 coding genes, with 132,970 coding transcripts. RNA-seq alignments suggested an average 19.5% (8-45%) improvement in completeness compared to Ensembl and RefSeq transcriptomes across all sequenced tissues (**Fig 8C**).

## 4 Discussion

Here we have presented a comprehensive equine transcriptome with tissue-specific expression utilizing the rich tissue repository from the FAANG biobank and advanced long-read sequencing technology. The FAANG equine transcriptome consists of 36,239 unique genes with 153,492 transcripts, presenting a gene-isoform ratio of 4.2, or 5.0 when only protein-coding genes are considered. This is a substantial improvement as compared to the ratios of 2.0 from the Ensembl equine transcriptome or 2.8 from the RefSeq equine transcriptome. We also demonstrated improved completeness of our transcriptome across over 40 tissues when assessed by companion RNA-seq data, despite having only 9 tissue types included in our Iso-seq experiment.

The previous efforts to annotate the horse genome were limited by the number of tissue types available and sequencing lengths available at that time^26–28^. Specifically, Hestand *et al*. sequenced 43 different equine tissues in one pool on an Illumina HiSeq 2000, both single-and paired-end at 75 bp on 4 lanes each^27^. While that study included more diverse tissue types than the current FAANG transcriptome, the pooling approach employed in that study limited discovery of rare novel and tissue-specific isoforms. The non-stranded protocol employed in that study also rendered it impossible to identify antisense transcripts. Mansour *et al*. compared sequences of 8 tissue samples from 59 individuals using short-read RNA-seq libraries from several studies (80-125bp, single-and paired-end, stranded and unstranded)^28^. Due to the limitation of short-read sequencing technology, in these studies, it was necessary to aggressively filter out mono-exonic transcripts that were not evolutionarily conserved, a common strategy in short-read based transcriptome assemblies^3,6^. This unfortunately would remove many small noncoding RNAs. Based on recent advances in long isoform sequencing, our approach centered around high-quality full-length reads from Iso-seq and used abundant RNA-seq data to refine splice junction, TSS, and TTS annotation. As a result, we were able to identify many novel mono-exonic transcripts as well as improve TTS annotation for many known transcripts.

In examining the transcriptional pattern of the horse genome, we revealed similar complexity in gene transcription to that of the human^7^. Specifically, we observed that genes with multiple isoforms tend to express more than one isoform simultaneously in any given tissue. The major isoforms (the isoform with highest expression) differed by tissue types. This aligns with the current understanding that isoforms, not genes, are directly associated with tissue-specific biofunctions. Most importantly, our data suggest that most known isoforms annotated in the Ensembl equine transcriptome are ubiquitous, while many novel isoforms identified in the Iso-seq transcriptome show tissue-specific expression. The addition of these novel isoforms should aid the equine genetics community in advancing studies of complex traits.

Despite these improvements, the present Iso-seq transcriptome was unable to accurately define TSS due to a lack of 5’ captured reads. Furthermore, small RNAs with no polyadenylated tails are missing from the poly-A captured cDNA libraries used for both Iso-seq and RNA-seq. Assays targeting non-polyadenylated RNAs such as small RNA sequencing and capturing 5’ capped transcripts like CAGE-seq are necessary to complement this Iso-seq transcriptome to fully capture the transcriptional landscape in the horse genome. Further, while we demonstrated improved completeness of the FAANG equine transcriptome, only 9 tissues were utilized to construct it, and many rare or tissue-specific transcripts are likely to be missing, especially stem-cell-specific or embryonically specific transcripts. Indeed, bone marrow was the only tissue that showed a drastic decrease in mapping rate when compared to RefSeq or Ensembl transcriptomes. These critical developmental time point and stem cell populations are required to further refine the horse transcriptome.

For the purpose of providing a comprehensive transcriptome, we focused on assessing the completeness of the FAANG equine transcriptome and overall complexities of tissue-specific transcription in the horse. However, the long read-length of Iso-seq data also provides unique opportunities for phasing the exons and, when coupled with whole genome sequencing and quantifiable RNA-seq of the same animals, studies of allele-specific expression. This is an area of future research. Additionally, the large repository of RNA-seq data from a diverse set of tissues enables studies of differentially expressed genes across tissues. Owing to the addition of the FAANG transcriptome, future studies can focus on quantifying gene expression across tissues and conditions.

## 5 Methods and Materials

### 5.1 RNA Extraction and Sequencing

Since the generation of the FAANG biobank, researchers were invited to “adopt” tissues of their interests, which meant they would sponsor the sequencing costs for two biological replicates (2 male or 2 female) of the “adopted” tissue. Under this Adopt-A-Tissue model, along with the eight prioritized tissues funded by USDA National Institute of Food and Agriculture and the Grayson Jockey Club Foundation, the equine community collectively generated short-read mRNA-seq data from over forty tissues. All RNA extractions for mRNA-seq were performed at the same two locations (female samples at UC Davis, male samples at University of Nebraska-Lincoln). Briefly, tissue aliquots were homogenized using Biopulverisor and Genogrinder in TRIzol reagent (ThermoFisher Scientific, Waltham MA). RNA was isolated and purified using RNeasy^®^Plus Mini/Micro columns (Qiagen, Germantown, MD) or Direct-zol RNA Miniprep Plus (Zymo Research, Irvine, CA). A detailed protocol can be found in **Supplemental Materials I and II**. For the female tissues, cDNA libraries were prepared with Illumina TruSeq Stranded kit and sequenced at the University of Minnesota sequencing core facility on an Illumina HiSeq 2500 using 125 bp paired-end reads. Male samples went through similar library preparation before 150 bp paired-end sequencing at Admera Health (South Plainfield, NJ) on an Illumina NovaSeq.

A total of nine tissues (lamina, liver, left lung, left ventricle of heart, longissimus muscle, skin, parietal cortex, testis, and ovary) from the FAANG biobank^17,18^ were selected for Iso-seq to represent a wide range of biological functions and therefore, gene expression. RNA for Iso-seq was extracted separately from the same tissues as mRNA-seq using the same protocol. All tissues for Iso-seq were processed in one batch. One sample per sex per tissue was selected for sequencing based on sample availability and RNA integrity numbers (RINs selected > 7). cDNA libraries were prepared and sequenced at UC Berkely QB3 Genomics core facility. Two libraries were randomly pooled and sequenced on a single SMRT cell on PacBio Sequel II.

### 5.2 Transcriptome Assembly

Pooled subreads were first demultiplexed using lima^29^. Circular consensus reads (ccs) were then constructed from demultiplexed subreads using PacBio ccs program^30^. PolyA tails were trimmed from ccs reads using isoseq3^31^. This step also removes concatemers and any reads lacking at least 20 bp of polyA tails. Redundant reads were then clustered based on pair-wise alignment using isoseq3^31^. Clustered transcripts were aligned to the reference genome EquCab3^2^ using minimap2^32^ without reference annotation as guide. Collapsed transcripts were filtered if they were not supported by at least two full length reads. Filtered transcripts from each sample were then merged into a single transcriptome using Cupcake^33^ and further filtered to retain only those detected in more than one sample. The merged total transcriptome was again aligned to the reference genome and collapsed to remove redundant transcripts. SQANTI3^23^ was then used to classify and annotate the transcriptome. Finally, the total transcriptome was filtered to remove nonsense-mediated decay transcripts, transcripts without short-read coverage support, and transcripts with a splice junction not covered by short-read RNA-seq data to generate the final FAANG equine transcriptome. Data processing, visualization, and statistical analyses were performed using pandas^34^, matplotlib^35^, seaborn^36^, scipy^37^, and scikit-learn^38^.

### 5.3 RNA-seq analysis

Short-read RNA-seq data were trimmed to remove adapters and low quality reads using trim-galore^39^ and Cutadapt^40^. Read qualities were inspected using fastQC^41^ and multiQC^42^. Trimmed reads were aligned to equCab3.0 using STAR aligner^43^ with standard parameters (with -- outSAMstrandField intronMotif --outSAMattrIHstart 0). PCR duplicates were marked using sambamba^44^. Mapping rates, qualities, and fragment lengths were assessed with samtools^45^ and deeptools^46^. Aligned reads were used to assess completeness of transcriptomes using deeptools^46^. BWA MEM^47^ was used to align the RNA-seq reads directly to transcriptomes and samtools^45^ was used to calculate the percentages of properly paired reads from the transcriptome alignment. Due to the presence of alternatively spliced isoforms in transcriptomes, multiple-alignment reads were not removed.

### 5.4 ATAC-seq analysis

ATAC-seq data from the 8 tissues (lamina, liver, left lung, left ventricle of heart, longissimus muscle, parietal cortex, testis, and ovary) collected from the same animals were generated and processed according to Peng *et al*.^21^ Libraries were sequenced in 50 bp paired-end mode (PE50) on Illumina NovaSeq 6000. Aligned reads were used to quantify normalized read counts in 1Kb up- and down-stream of TSS sites annotated in the equine FAANG transcriptome.

## 6 Data Access

RNA-seq data can be accessed from ENA and SRA under the accession number PRJEB26787. Iso-seq data can be accessed from ENA and SRA under the accession number PRJEB53020. ATAC-seq data can be accessed from ENA and SRA under the accession number PRJEB53037. Total and tissue-specific transcriptomes can be downloaded in FASTA and GTF format from https://equinegenomics.uky.edu/

## 7 Acknowledgements

The authors would like to acknowledge the four animals from which the FAANG tissues were collected.

