## Supplementary Material II for "Long-read RNA Sequencing Improves the Annotation of the Equine Transcriptome"

**Finno “Sesamoid modifications” of the RNeasy Lipid Tissue Kit (Qiagen)**

**Modified by Dr. Erin Hales for FAANG Isolations**

(<http://www.qiagen.com/products/rnastabilizationpurification/rneasysystem/rneasylipidtissuemini.aspx>)

- Have liquid nitrogen and dry ice ready
- Have cartilage, bone, or tendon frozen in liquid N_2_ as well as genogrinder plates cooled
- Have centrifuge at 4^O^C
- Have Qiagen micro columns (Cat No: C1004-50 from Direct-zol™ RNA MicroPrep kit)
- Warm RNase free water to 65^O^C
- Thaw Buffer RDD for DNase Digestion

1. Weigh 200 mg of bone pieces into a 5mL polyethylene spex mill tube and add a 9mm grinding ball. Freeze in liquid nitrogen in correct genogrinder trays.
2. Run the genogrinder for two 1-minute intervals at 1000 rpm. Do not look at the samples in-between. Just push the green button again! (Resist the dark side).
3. Freeze the samples and trays in dry ice for 1 minute and then liquid nitrogen again. Genogrind for 1 minute at 1200 rpm. Repeat this step once.
4. Freeze samples and trays one last time using both the dry ice and liquid nitrogen and run the genogrinder for 1 minute at 1500 rpm. Remove the samples and place in liquid nitrogen.
5. Remove the grinding ball using a magnet and Homogenize in the Spex Tube

Homogenization:

6. Add **1000 µl** **Trizol**® to pulverized tissue
7. Homogenize for 30/35s 3 times, 1min break between on ice all the time.
8. Add **1000 µl Trizol** so total volume is 2ml. Mix with pipette tip.
9. Cut the tip of a 1000ul tip and use it tip to move the homogenized sample into 15ml conical. Wash the homogenization tube with **2 mL Trizol**® and add to the 15 mL vial - **incubate** at room temperature for 30 minutes on a rotisserie light protected with a glove.
10. **Split** the cartilage-Trizol® homogenate into four 1.5 ml microcentrifuge tubes as 750ul/1.5ml.
11. Spin the tubes 10 minutes 12,000 x g 4^O^C
12. Transfer cleared supernatant to a clean 1.5 mL microcentrifuge tube
13. Add **200 μl chloroform** to each tube and Vortex 15 seconds. It will look like Pepto-Bismol®.
14. Incubate at room temperature 3 minutes.
15. Centrifuge 12,000 x *g* for 15 minutes at 4^O^C

RNA Collection:

16. **Transfer** aqueous phase (top layer) to a new 1.5ml tube. 200μl at a time.

- SC collection volume is ~500ul usually

17. **Add** one volume of **70% ethanol** (~500 μl but use tube markings) – mix by inverting 3 times.
18. Pipette **700 μl onto the RNeasy Micro Spin Column** in round bottomed collection. Centrifuge at **8,000 x *g* for 20 seconds** at 4^O^C. Discard the flow through.
19. Start water heating up.
20. **Repeat** step 12 until finish loading all volume of aqueous phase from the eight 1.5ml tubes
    a. Maintain all precipitate tubes on ice until ready for loading.
    b. Leave centrifuge open between spins so it can start warming to room temp for elution.

DNA Digestion:

21. Add **350 µl Buffer RW1**, centrifuge at **8,000 x *g* for 20 seconds** at 4^O^C. Discard flow through.
22. Add **70µl Buffer RDD** to a **10 µl Qiagen DNase aliquot** (found in the freezer). Mix by pipetting and add the **80 µl Reaction mixture** to the column.
23. Incubate at room temp for 15 minutes.

RNA Clean up and Collection:

24. Add **350 µl Buffer RW1,** centrifuge at 8,000 x *g* for 20 seconds at room temperature. Discard flow through. (**350 µl Buffer RW1** if DNA digestion was done.)
25. Add **500 μl Buffer RPE**, centrifuge at 8,000 x *g* for 20 seconds at room temperature. Discard flow through.
26. Add **500 μl Buffer RPE again** (for micro kit, this step is 80% ethanol), centrifuge at 8,000 x *g* for **2 minutes** at room temperature. Discard flow through and the collection tube.
27. Place Spin Column in new collection tube. Open lid to dry; centrifuge for **5 minutes at full speed**. Discard flow through and the collection tube.
28. Place spin column in a new, labeled tube, add **20 μl RNase-free** **water** (65^O^C). Incubate at room temperature for 60 seconds. Spin at **8,000 x *g* for 1 minute** at room temperature.
29. Run Nanodrop and **store at -80^o^C**. Run Bioanalyzer when enough samples have been isolated.

Protocol adapted from Annette Marie of the McCoy lab and Jamie MacLeod lab September 2017
