## Supplementary Material I for "Long-read RNA Sequencing Improves the Annotation of the Equine Transcriptome"

**RNA Isolation Using a Column and on-column DNA digestion**

Preparation:

1. Wash area around balance and tools with RNase away.
2. Disrupt ~70 mg of sample tissue (100 mg of adipose tissues) on dry ice.
3. Place minced aliquot into a 2.0 ml tube and place in -80^O^C freezer.
4. Label a set of columns (Zymo Research Direct-zol RNA Miniprep kit, Irvine, CA, USA), 1.5 ml tubes, and RNA final elution tubes.
5. Set aside a labeled waste beaker for use during the wash steps.

Tissue Homogenization:

1. Wash all pipettes and work benches with RNase away.
2. Add **600 µl Trizol** (Invitrogen, Waltham, MA, USA) to 2.0 ml tube containing pre-weighed, minced tissue.
3. Disrupt tissue using homogenizer at appropriate speed for specific tissue type in 30 second bursts ON ICE. (Homogenize for 30 seconds and then let sample rest on ice for 30 seconds) Check homogenization by using needle and syringe and pipetting up and down. Add **400 µl Trizol** and mix by pipetting.
   a. Keep homogenized samples on ice while homogenizing other samples.
4. Incubate all samples for 5 minutes at room temperature. (Critical)
   a. Tough tissues including bone, cartilage, tendon, adipose, and cornea require a longer benchtop incubation sometimes up to 30 minutes.
5. For difficult tissues, centrifuge at 12,000 x g for 10 minutes at 4°C and transfer cleared, pink supernatant to a new labeled tube avoiding cellular debris and uppermost clear fat layer.
   a. This includes adipose, skin, cornea, cartilage and bone
6. Add **200** **µl** **chloroform** and vortex for 15 seconds, then incubate at room temperature for 2-3 minutes.
7. Centrifuge at 12,000 x g for 15 minutes at 4°C.
8. Add **600 µl 100 % ethanol** to a set of labeled 1.5 ml tubes while the centrifuge is running.

RNA Collection:

1. Transfer clear aqueous supernatant to the appropriate labeled ethanol tubes 200 µl at a time. Be very careful to not pipette up any of the pink or white layers. It will make the final RNA more “dirty” (i.e. Decreasing the ratios and RIN score).
2. Invert tubes until solution is homogenous (~3 inversions).
3. Load 600 µl of the ethanol mixture onto the appropriately labeled column.
4. Centrifuge for 30 seconds at 12,000 x g.
5. Discard flow through into waste beaker and load remaining ethanol mixture onto the column. Centrifuge for 30 seconds at 12,000 x g. Discard flow through.

DNA Digestion:

1. Wash the column with **400 µl RNA Wash Buffer**. Centrifuge for 1 minute at 12,000 x g; discard flow through.
2. Prepare **DNase I Reaction Mix** for each sample by adding **75 µl DNA digestion Buffer** to a **5 µl aliquot of DNase I**. *These aliquots are stored at -20^o^C. *
3. Add the **80 µl** **DNase I Reaction Mix** directly to the column.
4. Incubate at room temperature for 15 minutes.

RNA Collection:

1. Add **400 µl** **Direct-zol RNA PreWash** to the column then centrifuge at 12,000 x g for 1 minute. Discard the flow through. **Repeat** this step once.
2. Add **700 µl** **RNA Wash Buffer** to the column and centrifuge at 12,000 x g for 1 minute. Discard flow through and spin again for 2 minutes at 12,000 x g.
3. Transfer the column to the labeled RNA final elution tube.
4. Add **50-100 µl** **DNase/RNase-free water** to the column and load into centrifuge with caps pointed towards the lower numbers (the right). This is critical to prevent the caps from being snapped off during the centrifuge cycle.
5. Centrifuge at 12,000 x g for 1 minute
6. Spec the RNA by loading 2 µl per sample into a slide and running it on the QIAxpert. Make sure this pipette is cleaned before you use it. *If you do not plan on spec’ing the samples immediately, you may aliquot 2 µl in a separate labeled tube and freeze. This will help limit the number of freeze-thaw cylces.
7. Store RNA at -80^O^C.
